# link-ancestors: Fast simulation of local ancestry with tree sequence software

**DOI:** 10.1101/2023.11.03.564476

**Authors:** Georgia Tsambos, Jerome Kelleher, Peter Ralph, Stephen Leslie, Damjan Vukcevic

**Author notes:** All of these authors contributed equally (Jerome Kelleher, Peter Ralph, Stephen Leslie, Damjan Vukcevic).

## Abstract

**Summary:** It is challenging to simulate realistic tracts of genetic ancestry on a scale suitable for simulation based inference. We present an algorithm that enables this information to be extracted efficiently from tree sequences produced by simulations run with msprime and SLiM.

**Availability and Implementation:** A C-based implementation of the link-ancestors algorithm is in tskit (https://tskit.dev/tskit/docs/stable/). We also provide a user-friendly wrapper for link-ancestors in tspop, a Python-based utility package.

**Supplementary information:** Technical details of link-ancestors are in the Supplementary Information. Documentation for the tspop package is at http://tspop.readthedocs.io/.

## 1 Introduction

One way to think about a sample’s genetic ancestry is to consider all of the genetic ancestors who existed at some specific time point in the past. When analysing or visualising this ancestry, it is often convenient to partition the ancestors into groups and to “paint” each part of each sampled chromosome according to their *local ancestry*, i.e., which group it was inherited from. In practice, these distinct groups are usually called “ancestry components” or “populations” (Coop, 2022), and conceptually correspond to relatively distinct historical groups, often partitioned by geography. An understanding of genetic ancestry is central to the application and interpretation of many methods in population genetics, medical genetics and bioinformatics. Accordingly, the ability to describe it in simulations is valuable. While dedicated local ancestry simulators exist (Corbett-Detig and Jones, 2016; Schumer *et al*., 2020), they are limited in the range of evolutionary processes that they model, and they do not simulate genetic variation. The popular msprime and SLiM programs allow users to simulate large genomes and sample sizes efficiently under a wide range of evolutionary processes, demographic models and genomic parameters (Kelleher *et al*., 2016; Baumdicker *et al*., 2022; Haller and Messer, 2018, 2023). Moreover, they are capable of outputting complete genetic genealogies in a compact data format, the succinct tree sequence (Kelleher *et al*., 2016). Although these simulated outputs contain great detail about genomic ancestry, extracting this information is a difficult problem. This note describes link-ancestors, an algorithm that takes a simulated tree sequence and a set of ancestors, and efficiently returns a record of the corresponding local ancestry. We present two tools: a low-level function in the open-source tskit package, and a small utility package, tspop, that provides more user-friendly output for msprime and SLiM users.

## 2 Materials and Methods

The succinct tree sequence data structure was introduced by Kelleher *et al*. (2016) as a way to represent simulated gene genealogies compactly. All tree sequences contain the following minimal ingredients: a set of nodes N, which represent contemporary and ancestral haplotypes of the genomic sample; and a set of edges E, which represent lines of descent between the nodes. Each node n ∈ N is associated with a time tn. Each edge e ∈ E is associated with a parent node pe ∈ N, a child node ce ∈ N such that tce < tpe, and left and right coordinates [le, re). This indicates that the child ce has inherited the genomic segment [le, re) from parent pe. Because the trees contain the full genomic history of the simulated sample, they are considerably more informative than raw haplotype data, and they can also be queried and modified quickly, enabling rapid computation of statistics across the genome (Ralph *et al*., 2020).

Unlike dedicated local ancestry simulators (Corbett-Detig and Jones, 2016; Schumer *et al*., 2020), which only simulate information about ancestral tracts, the output from tree sequence simulators like msprime and SLiM can in theory be queried for any site-or ancestry-based information in the genotypes or gene genealogies. These properties make them useful for simulation-based inference procedures, such as approximate Bayesian computation, where one must typically compute many complementary summary statistics on each simulated dataset, as well as in exploratory simulations, when one is not certain which properties of the simulated genomes will be of most interest ahead of time. An additional advantage of tree sequence based software is its considerable flexibility and modular inter-operability. Users can run rapid backwards-in-time simulations with msprime or ultra-realistic forwards-time simulations with SLiM, combine them with pyslim (Ralph *et al*., 2022), and analyse them with tskit (Baumdicker *et al*., 2022; Haller and Messer, 2018; Ralph *et al*., 2020). Our addition of link-ancestors into tskit extends its utility to a range of haplotype-based analyses.

Technical details of the link-ancestors algorithm are included as a supplement, but we provide some intuition here. The user must input a tree sequence, typically simulated using msprime or SLiM under a model of population structure, as well as a set of ancestors from the ancestral populations of interest (Figure 1A). To ensure that local ancestry is well defined with respect to a specific time, we recommend setting up the simulation so that all ancestors at a specific ‘census time’ are explicitly recorded and passed to link_ancestors(). This can be done using a census_time() event in msprime, or with a treeSeqRememberIndividuals() callback in SLiM. Each edge in the tree sequence represents a segment of genome that has been inherited by a particular descendant from a particular ancestor (Figure 1B), and at each point in the past, there is a set of segments in the ancestral genomes that each modern day genome has inherited from. We can therefore identify those segments by tracing through the relationships described by the edges.

**Fig. 1.**
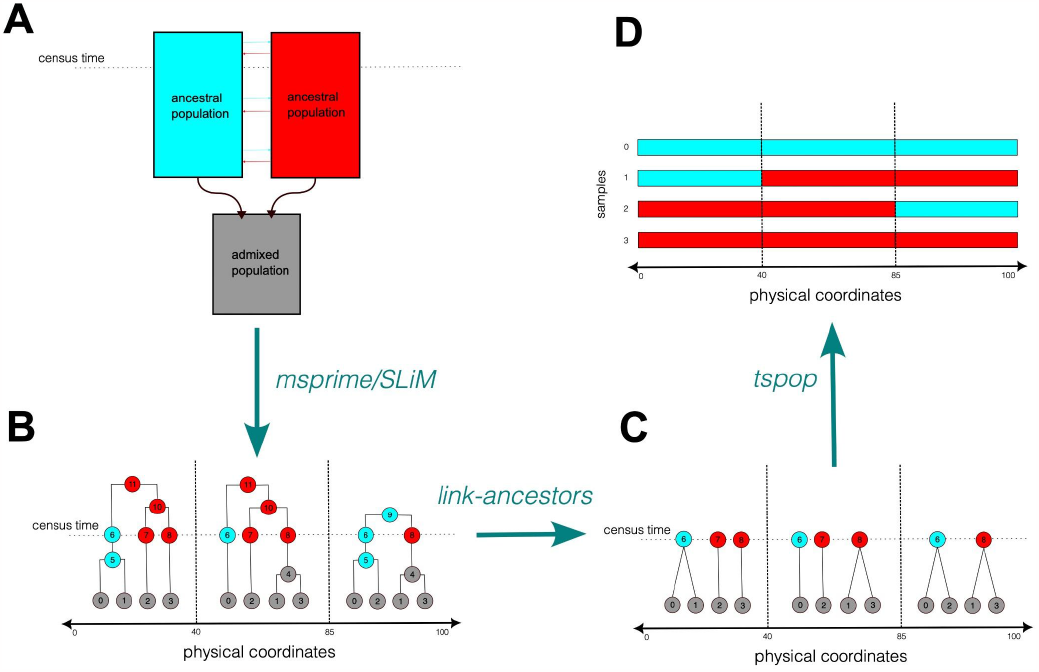
A visualisation of the pipeline that can be used to simulate local ancestry with link-ancestors. (A) An admixture model that can be programmed with msprime and SLiM, and from which simulated genomes can be drawn. In this simple example, individuals in a red and a blue population with small amounts of gene flow then undergo admixture, and all subsequent genomes belong to individuals in an admixed grey population. A census event prior to the admixture indicates the time of the ancestors that local ancestry will be defined with respect to. (B) A sample tree sequence that may be outputted by the simulation run in step A. Six admixed genomes of length 300 bp are labelled 0-5. Ancestral genomes are coloured according to the population of the individual they belong to. The breakpoints between the trees indicate locations of past recombination events in the sample. Note that an ancestral genome is recorded on each genomic lineage that is extant at the time of the user-specified census. (C) A visualisation of the output produced by the link-ancestors method when applied to the tree sequence shown in (B). All edges on paths joining a sample genome to a census genome have been concatenated into a single edge between the two genomes. (D) The admixture tracts of the sample genomes shown in (C). These can be obtained by replacing the label of each census genome in (C) with the population it belongs to, a function that is provided in the tspop package. Note that these admixture tracts depend on the timing of the census event; (B) shows that, for instance, samples 1 and 3 have ancestry with the blue population between coordinates 85 and 100 when viewed with reference to the time of ancestor 9.

The algorithm works by iterating through the edges, which are ordered by time, until the labeled ancestors are reached. At each step, the algorithm records the ancestral segments from which each focal genome has inherited as its internal state. These records are updated using the information in each successive edge; therefore, as the algorithm moves through edges representing older ancestors, segments are divided into smaller and smaller pieces. The output is an edge table corresponding to a smaller, simplified tree sequence (Figure 1C). Each edge in this output shows the coordinates of some tract of local ancestry that a sample has inherited from one of the ancestors in the user-specified groups of interest. However, often only the ancestor’s population label is of interest. For this use case, the tspop package contains additional functions to ‘squash’ these small segments into longer contiguous segments from the user-specified population labels, and to perform simple analyses using this condensed output (Figure 1D).

Performing this operation separately on each local tree in the gene genealogy is inefficient; in general, many edges are shared due to recent common ancestry between the samples. By storing the common parts of these calculations in memory, link-ancestors works in a single pass over the set of edges, greatly reducing the total number of operations. We compare link-ancestors with this tree-by-tree approach in the Supplementary Information.

## 3 Usage

The efficient C-based implementation of the link-ancestors algorithm described here is contained in the tskit package (Ralph *et al*., 2020), and can be called from a wrapper in the user-facing Python API. Users input a tree sequence along with a list of sample and ancestral nodes. The raw output of the link-ancestors function is an edge table satisfying the conventions of the tree sequence format (Kelleher *et al*., 2016). Each row is of the form (*l, r, a, c*), indicating that sample node *c* is descended from ancestral node *a* on the interval [*l, r*).

The tspop package contains a wrapper function for link-ancestors that also performs a number of other post-processing operations that one typically requires in a local ancestry-based analysis. tspop can annotate link-ancestors output with an extra column, *p*_*a*_, indicating the population that *a* belonged to. They are provided as reordered pandas dataframes (Reback *et al*., 2022) for ease of downstream analyses. tspop can also summarise and visualise this output in various ways.

To demonstrate the performance of these methods, we used link-ancestors to identify segments of continental ancestry dating to 100 generations in the past, in datasets that had been simulated to resemble chromosome 1 in 1000 modern-day admixed American human individuals. This took around 4 sec using around 0.5 Gb of RAM.

Note that link-ancestors stores genomic segments in memory at every node along each path connecting a sample to a census ancestor of interest. Nodes representing older ancestors will generally be ancestral to a larger number of descendant samples, and will thus require more genomic segments to be stored. Because of these factors, link-ancestors requires more RAM to extract segments of local ancestry from older ancestors, or from simulations with a larger number of ancestors that are explicitly recorded in the trees (as in a pedigree-based simulation, for instance). This can come at a cost of increased memory usage. To illustrate this, we ran another simulation designed to model archaic introgression approximately 2000 generations ago across chromosome 1 in the ancestry of around 100 modern humans. Applying link-ancestors to identify ancestry from the time of introgression from this simulation requires around 61 seconds and 9 Gb of RAM.

Assuming a per-base, per-generation recombination rate of ϱ and a sample chromosome of physical length *l*, we expect a total of ϱ*l* new recombination breakpoints separating the chromosome from its genetic ancestors in the prior generation. In each successive generation backwards-in-time, this ancestral material may be spread between more ancestors, but the total length of these ancestral segments is unchanged. Therefore, each generation is expected to add ϱ*l* to the total number of distinct segments that the modern genome has inherited from. This means that the expected number of distinct segments that a given modern genome inherits from at a given time in the past scales linearly with the amount of time. For this reason, we expect the size of the output to scale linearly with the time between the samples and ancestors, as well as with the number of samples and the length of the genome. However, since the algorithm keeps track of ancestral segments at intermediate times as well, the required memory will scale faster than linearly with time (backwards). Given ancestors in a specific epoch of time, the total number of algorithmic steps scales according to the size of the inputted edge table (see Kelleher *et al*., 2016).

Users should also be aware that the outputted local ancestry tracts may depend on the specific choice of ancestors supplied to the link-ancestors algorithm, especially if the simulation involves ongoing migration between the ancestral populations of interest. Consider a genomic ancestor *a*□ who belonged to a certain population *n* generations ago, and whose child *a*□_−1_ at time *n*-1 generations ago migrated into a different population. Any present-day genomic segments that descend from these two ancestors will have passed through both populations, so their local ancestry is ill-defined without any extra information. link-ancestors uses the user-specified set of ancestors to determine the assignment; if the set of provided ancestors is {*a*□, *a*□_−1_} (or just {*a*□_−1_}), the segment will be linked to the most recent ancestor *a*□_−1_, but if the set is {*a*□} it will be linked to the older ancestor *a*□. To avoid confusion, we suggest that population-based ancestry is defined with reference to ancestors at a single census timepoint, as described in the Materials and Methods.

## 4 Conclusion

Tools that can simulate genetic ancestry are crucial to many analyses. The link-ancestors algorithm performs the essential, difficult step of extracting local ancestry from a simulated tree sequence. A future improvement to our software would be to add a ‘garbage-collection’ procedure to improve memory usage on very deep timescales (Corbett Detig and Jones, 2016). We hope that our open-source implementations in tskit and tspop will facilitate exploration, benchmarking and inference, and in turn lead to new insights about population-based ancestry.

