## Supplementary Material for "link-ancestors: Fast simulation of local ancestry with tree sequence software"

#### The `link-ancestors` algorithm

Recall that all tree sequences contain the following minimal ingredients: a set of *nodes* $\mathcal{N}$, which represent contemporary and ancestral haplotypes of the genomic sample; and a set of *edges* $\mathcal{E}$, which represent lines of descent between the nodes. Each node $n \in \mathcal{N}$ is associated with a time $n_t$. Each edge $e \in E$ is associated with a parent node $p_e \in \mathcal{N}$, a child node $c_e \in \mathcal{N}$ such that $t_{c_e} < t_{p_e}$, and left and right coordinates $[l_e, r_e)$. This indicates that the child $c_e$ has inherited the genomic segment $[l_e, r_e)$ from parent $p_e$.

To extract local ancestry from the output described above, the tree sequence needs to have *census nodes*: nodes on all extant lineages at the ancestral time of interest. This can be done with the `add_census()` method in `msprime` or a `treeSeqRememberIndividuals()` call in `SLiM`. These ancestral nodes should also be labeled (we call the labels "populations" although other use cases are possible). Then, at each location along the simulated genome, trace a path through the edges to determine which ancestral nodes are ancestral to each sample node. If a sample node $s$ is descended from an ancestral node $a$ on the genomic interval $[l, r)$, then it has local ancestry with the population label of $a$ on that interval. Tracing these paths is potentially complicated and inefficient, and is the operation performed by `link-ancestors`.

This section presents a pseudo-coded set-theoretic description of the `link-ancestors` algorithm. The procedure is shown in Algorithm 1, and a reference of objects in this algorithm is in Table 1. It is a modification of the algorithm for tree sequence 'simplification' (Kelleher *et al.*, 2018). As input, the user must provide a tree sequence with nodes $\mathcal{N}$ and edges $\mathcal{E}$, a set of sample nodes $\mathcal{C} \subset \mathcal{N}$ and a set of ancestral nodes $\mathcal{P}$. See Kelleher *et al.* (2016) for more information about these tree sequence objects.

The outermost loop iterates over the rows of the edge table (line 2), which are sorted first by the time of the parent node. Lines 3 to 15 construct a set of sample segments $\mathcal{S}$ that descend from the parent node $p_i$ of the edge $e_i$ currently being processed. These are later used to update a master list $\mathcal{A}_{p_i}$ of segments descending from the node. Lines 16 to 21 show the steps that are invoked when the algorithm finds an edge with a parent node that belongs to the supplied list of ancestral nodes. At this point, each segment is used to construct an edge in the output table indicating its direct descent from the ancestral node (line 19).

**Algorithm 1** The `link-ancestors` algorithm.

Set $i \leftarrow 1$, $\mathcal{S} \leftarrow \emptyset$ and $\mathcal{A}_n \leftarrow \emptyset$ for all $n \in \mathcal{N}$.
**while** $i \leq E$ **do**
  **if** $c_i \in \mathcal{C}$ **then**
    $\mathcal{S} \leftarrow \mathcal{S} \cup \{ (c_i : [l_i, r_i)) \}$
  **else**
    **if** $|\mathcal{A}_{p_i}| > 0$ **then**
      $j \leftarrow 1$
      **while** $j \leq |\mathcal{S}|$ **do**
        $(c_j : [l_j, r_j)) \leftarrow \mathcal{A}_{p_i}(j)$
        $l \leftarrow \max\{l_i, l_j\}$, $r \leftarrow \min\{r_i, r_j\}$
        $\mathcal{S} \leftarrow \mathcal{S} \cup \{ (c_j : [l, r)) \}$
        $j \leftarrow j + 1$
      **end while**
    **end if**
  **end if**
  **if** $p_i \neq p_{i-1}$ or $i = E$ **then**
    **if** $p_i \in \mathcal{P}$ **then**
      **for** $(c_j : [l_j, r_j)) \in \mathcal{S}$ **do**
        $\mathcal{L} \leftarrow \mathcal{L} \cup (l_j, r_j : p_i, c_j)$
      **end for**
    **end if**
    $\mathcal{A}_{p_i} \leftarrow \mathcal{A}_{p_i} \cup \mathcal{S}$
    $\mathcal{S} \leftarrow \emptyset$
  **end if**
  $i \leftarrow i + 1$
**end while**

| Label | Object type | Description |
|---|---|---|
| $\mathcal{A}_n$ | Internal | A set of all segments descending from ancestor $n \in \mathcal{N}$ with a child node in the set of sample pairs. |
| $\mathcal{C}$ | User-specified | A list of sample nodes for which we wish to calculate local ancestry. |
| $c_i$ | Internal | The child node of the $i$th edge in $\mathcal{E}_{\mathcal{N}}$. |
| $E$ | Internal | The total number of edges in the edge table $\mathcal{E}_{\mathcal{N}}$. |
| $\mathcal{E}_{\mathcal{N}}$ | Tree sequence | The ordered set of edges in the tree sequence. |
| $i$ | Internal | An index for the edges in $\mathcal{E}_{\mathcal{N}}$. |
| $j$ | Internal | An index for the segments in $\mathcal{S}$. |
| $k$ | Internal | An index for the segments in $\mathcal{A}_n$. |
| $l_i$ | Internal | The left coordinate of the $i$th edge in $\mathcal{E}_{\mathcal{N}}$. |
| $m$ | Internal | An index for the segments in $\mathcal{S}$. |
| $\mathcal{L}$ | Output | The outputted edge table. |
| $\mathcal{N}$ | Tree sequence | The ordered set of nodes in the tree sequence. |
| $p_i$ | Internal | The parent node of the $i$th edge in $\mathcal{E}_{\mathcal{N}}$. |
| $r_i$ | Internal | The right coordinate of the $i$th edge in $\mathcal{E}_{\mathcal{N}}$. |
| $\mathcal{P}$ | User-specified | A list of ancestral nodes from which we wish to calculate local ancestry. |
| $\mathcal{S}$ | Internal | An ordered list of genomic segments of the form $\big(c : [l, r)\big)$. |

Table 1: A reference of objects in the `link-ancestors` algorithm.

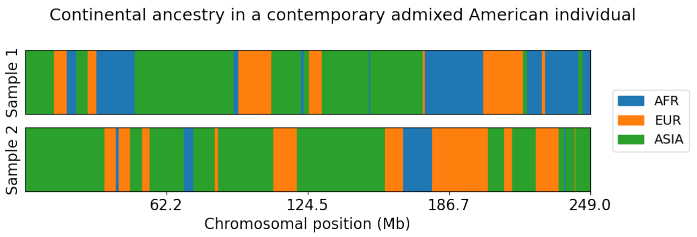


Figure 1: A pair of modern-day American human chromosomes are simulated with `msprime` to contain tracts of ancestry from three sources: Africa (AFR), Europe (EUR) and Asia (ASIA). This is an example of the information that can be extracted with the `tspop` wrapper for `link-ancestors`. See the Supplementary Information for more information about the simulation.

### Details of the American admixture exploratory simulation

Our first demonstration simulation uses the American admixture model originally published by Browning *et al.* (2018) and implemented in `stdpopsim` (Adrion *et al.*, 2020; Lauterbur *et al.*, 2022).

This model contains four modern-day populations representing contemporary *Homo sapiens*: AFR (Contemporary African population), EUR (Contemporary European population), ASIA (Contemporary Asian population) and ADMIX (Modern admixed population). The ADMIX population is modelled as an instantaneous pulse of gene flow from the other three populations 10 generations in the past. There is also a small amount of continual migration between the other three populations going back to 1000 generations in the past. See the `stdpopsim` catalog for full details of this model. We simulated a diploid region of length 243,199,373 base pairs with a uniform per-base, per-generation recombination rate of $1.15235 \times 10^{-8}$, as per the `stdpopsim` defaults for *Homo sapiens* chromosome 1.

To ensure the correctness of our model, we exported the `stdpopsim` implementation into `msprime` code using the Demes package (Gower *et al.*, 2022). We then added a census event at 1000 generations in the past. The ancestral nodes recorded at this census time were then used as input into `link-ancestors`. The choice of census time ensures that these ancestors belong to one of the three populations: AFR, EUR or ASIA.

We generated 10 replicate tree sequences holding 1000 individuals (2000 chromosomes) from each of the three contemporary populations. Tracts of local ancestry from these ancestors in one simulated present-day individual are plotted in Figure 1. The code we ran is available at `https://github.com/gtsambos/ooa-ancestry-example`.

| Approach | Runtime (sec) | RAM (Mb) |
|---|---|---|
| `link-ancestors` | $4.40 \pm 0.64$ | $535.86 \pm 0.13$ |
| 'baseline' | N/A | $> 128\,000$ |
| 'squash' | $21832.92 \pm 671.60$ | $487.29 \pm 0.13$ |
| 'numba' | $6136.87 \pm 891.62$ | $526.57 \pm 0.17$ |

Table 2: Means and standard deviations of runtimes and RAM usage from various approaches to extract local ancestry from tree sequences. The simulations used a demographic model of contemporary American human admixture using a census time of 100 generations in the past.

#### Details of the runtime and memory comparisons

On each of the replicates, we compared the performance of `link-ancestors` with several simpler tree-by-tree approaches using `tskit`'s tree methods. Runtimes were calculated using Python's `time` module, while memory usage was estimated using the `scalene` command-line interface (Berger *et al.*, 2020). Our results are shown in Table 2.

The initial approach that we considered, 'baseline', involved a loop through each tree $T$ and each sample node. At each sample node, we iterated upwards through the tree to obtain a local genealogical path. Once we encountered an ancestral node from the census time of 100, we terminated the iteration and recorded a tract of local ancestry between the sample and the ancestor over the interval spanned by the tree. This procedure requires of order $O(Tn\log(n))$ operations for a dataset of $n$ samples in a tree sequence of $T$ trees; since many segments of ancestry span multiple trees, this is much less efficient in practice than `link-ancestors`. However, this procedure had substantial memory requirements, requiring over 128 Gb of RAM. This is likely due to the uncompressed nature of the output: since consecutive trees differ by just one subtree-prune-and-regraft operation, many samples will share a common census ancestor across many consecutive trees. Therefore, instead of proceeding with this approach, we ran a modified version, 'squash', that progressively compressed these neighbouring segments of identical ancestry. We also considered that some of the runtime differences were due to the fact that the computationally intensive part of our `link-ancestors` implementation is coded in C. To minimise the impact of this difference, we used the `numba` package to create a streamlined pre-compiled version of our 'squash' implementation; we refer to this as the 'numba' procedure. The 'numba' procedure was substantially faster than 'squash' on its own, and with a minimal cost to memory usage. However, both methods required several hours of computation, and were thus still orders of magnitudes slower than the implementation of `link-ancestors` in `tskit`, which ran in roughly 4 seconds. Differences in memory requirements between these three methods were negligible.

| Approach | Runtime (sec) | RAM (Gb) |
|---|---|---|
| `link-ancestors` | $59.57 \pm 15.68$ | $9.02 \pm 2.05$ |
| ‘baseline’ | $1885.52 \pm 152.90$ | $78.76 \pm 0.00$ |
| ‘squash’ | $3705.63 \pm 671.60$ | $0.65 \pm 0.00$ |
| ‘numba’ | $1140.34 \pm 122.59$ | $0.83 \pm 0.00$ |

Table 3: Means and standard deviations of runtimes and RAM usage from various approaches to extract local ancestry from tree sequences. The simulations used a demographic model of Neanderthal and other archaic human introgression using a census time of 2093.5 generations in the past.

#### Details of the Out-of-Africa exploratory simulation

To highlight how the performance of `link-ancestors` varies according to the time of the census ancestors, we briefly describe some simulations using an older census time. We chose the Out-of-Africa model with archaic admixture originally published by Ragsdale and Gravel (2019) and also implemented in `stdpopsim` (Adrion *et al.*, 2020; Lauterbur *et al.*, 2022).

This model contains three modern-day populations representing contemporary *Homo sapiens*: YRI (Yoruba, African), CEU (European, Utah) and CHB (Han Chinese, Beijing). There are two populations representing archaic hominins: NEA (Neanderthals) and ARC (a ‘ghost’ archaic African population). Under this model, there are small amounts of ancient introgression between the following pairs of populations: (YRI, ARC), (CEU, NEA), (CHB, NEA). There is also migration between the three *Homo sapiens* populations. The ‘out-of-Africa’ event is modelled as an instantaneous split within the YRI population 2093 generations in the past. See the `stdpopsim` catalog for full details of this model. The simulated contig was the same as before. We added a census event at 2093.5 generations in the past, just prior to the Out-of-Africa event in which the CEU and CHB lineages split from YRI. This choice of time ensures that the census ancestors belong to one of three populations: YRI, NEA and ARC. We generated 10 replicate tree sequences holding 100 individuals (200 chromosomes) from each of the three contemporary populations. (The smaller sample size was chosen to ensure that we could test `link-ancestors` on a local machine with <16 Gb of RAM).

Results of these tests are shown in Table 3. Although `link-ancestors` is still an order of magnitude faster the other approaches, the relative difference is less striking than in the American admixture example detailed in the previous section. It also requires a larger amount of memory. This is due to the fact that the tree-by-tree methods do not need to retain any information about ancestors in previous trees, unlike `link-ancestors`. We therefore expect that `link-ancestors` will prove most useful in settings where the admixture events of interest are not ancient.
